# CRIMP: A CRISPR/Cas9 Insertional Mutagenesis Protocol and the CRIMP Toolkit

**DOI:** 10.1101/2023.07.13.548929

**Authors:** Lee B Miles, Vanessa Calcinotto, Carmen Sonntag, Clara Lee, Robert J Bryson-Richardson

## Abstract

We developed a highly efficient targeted insertional mutagenesis system, CRIMP, and an associated plasmid toolkit, CRIMPkit, that disrupts native gene expression by inducing complete transcriptional termination to produce null mutant alleles without inducing genetic compensation. The CRIMPkit contains over 30 ready-to-use plasmid vectors allowing easy and complete mutagenesis of any gene in any reading frame without requiring custom sequences, modification, or subcloning, and also provides a fluorescent readout of successfully mutagenised fish.

---

CRISPR/Cas9 technology has become a widely used tool for genome editing in model organisms, transforming functional genetics. Challenges, however, do remain. The introduction of nonsense mutations from the insertion of stop cassettes or generation of indel mutations has the potential to induce genetic compensation (Rossi et al., 2015; Sztal et al., 2018; El-Brolosy et al., 2019), masking loss of function phenotypes. Additionally, the efficiency of editing can vary dramatically depending on the complexity of the modification, and therefore substantial time and cost can be involved in genotyping and identifying founders. Targeted integration has the potential to overcome both these issues, and site-directed integration of foreign sequence through homology directed repair (HDR) has been successful, with inserts ranging from short DNA oligonucleotides (Gagnon et al., 2014) to entire expression cassettes (Welker et al., 2021; Levic et al., 2021; Gratz et al., 2014). However, several drawbacks, such as requiring a custom synthesis of a targeting vector for each target gene, and low integration efficiencies, have limited the use of HDR. Targeted integration via non-homologous end-joining (NHEJ) is more efficient than HDR but more error-prone, making it less appealing for targeting coding sequences. However, an intron-targeting strategy where a splice acceptor and downstream coding sequence are inserted into the intronic region of a gene truncating the protein overcomes this issue and has been used to generate mutant alleles (Kesavan et al., 2018; Li et al., 2019; Han et al., 2021; Lu et al., 2021). Although more efficient than HDR, the NHEJ-intron targeting approach has previously required individual targeting vectors to be cloned for each gene, limiting its broad application.

To overcome these issues and enable a broader uptake of NHEJ mediated insertional mutagenesis we have developed a universal and highly efficient targeted insertional mutagenesis system and toolkit that disrupts native gene expression by inducing complete transcriptional termination to produce full mutagenesis without genetic compensation and demonstrate its use in zebrafish.

The CRISPR/Cas9 Insertional Mutagenesis Protocol (CRIMP) and toolkit (CRIMPkit) contains 30 ready-to-use universal plasmid vectors to allow easy and complete mutagenesis in any reading frame without requiring custom sequences, modification, or subcloning. All CRIMPkit vectors are available from Addgene (199469-199498). Once inserted, these vectors drive a fluorescent reporter under the control of the targeted gene promoter, disrupting native gene expression and providing a reporter of the targeted genes expressions pattern and visual identification of successfully mutagenised fish. The toolkit contains multiple versions of each plasmid with different fluorophores; *mTagBFP2, mKate2*, and the *splitGFP* system to visually genotype fish and facilitate automated high-throughput screening, as well as multiple reporter expression systems enabling visual selection of fluorophores when expressed from both high to low expressed target genes. The plasmid vector design is outlined in Supplementary Figure S1, and the complete list of plasmids is provided in Table S1.

A challenge of site-directed insertion is the low efficiency of integration events, requiring many animals and extensive screening to identify founder animals. Our optimised protocol (Methods, Supplementary Figure S2) enables highly efficient integration, as evidenced by mosaic transgene expression in the majority of injected embryos. Furthermore, in a small number of embryos, the correct reporter expression pattern could be detected in exactly half of the embryo body plan (Supplementary Figure S3), indicating that integration occurred rapidly after injection, either during or directly after the first cell division. In our experience, all animals showing this pattern successfully passed on the insertion to the next generation. This enables preselection of founders prior to raising, reducing the number of animals raised, as well as the time and cost to screen founders. Where this pattern was not identified, we raised fish showing mosaic fluorescent transgene expression and were also able to successfully identify founders amongst these fish.

To demonstrate the ability of CRIMPkit vectors to generate mutant alleles, we selected *tdgf1* as a target gene, as it has a very well-characterised mutant phenotype (*one-eyed pinhead*, Schier et al., 1996; Solnica-Krezel et al., 1996; Zhang et al., 1998; Ata et al., 2018). We inserted the *pSA2-T2A-Gal4vp16/4xnrUAS-mTagBFP2* vector into intron-3 of *tdgf1*, to generate the targeted insertion line *Ti(tdgf1*^*int3*^*-Gal4vp16/4xnrUAS-mTagBFP2)* (Figure 1A, and Supplementary Figure S4). 95% of injected embryos had detectable mosaic *mTagBFP2* expression, demonstrating high efficiency, and included one embryo displaying the correct expression pattern throughout the ventral half of the body plan (0.7% of injected embryos). When raised to adulthood and crossed to wildtype this individual passed on the transgenic insertion, successfully establishing a *tdgf1* mutant line (Supplementary Figure S4). Heterozygous carriers are phenotypically wildtype with *mTagBFP2* expressed in the same pattern as the published expression data for *tdgf1* (Zhang et al., 1998). Embryos homozygous for the insertion phenocopy *tdgf1* mutants displaying eye and head defects and a ventrally curved body (Figure 1A) as previously reported (Schier et al., 1996; Solnica-Krezel et al., 1996; Zhang et al., 1998), demonstrating the ability of CRIMPkit vectors to generate mutant alleles.

**Figure 1.**
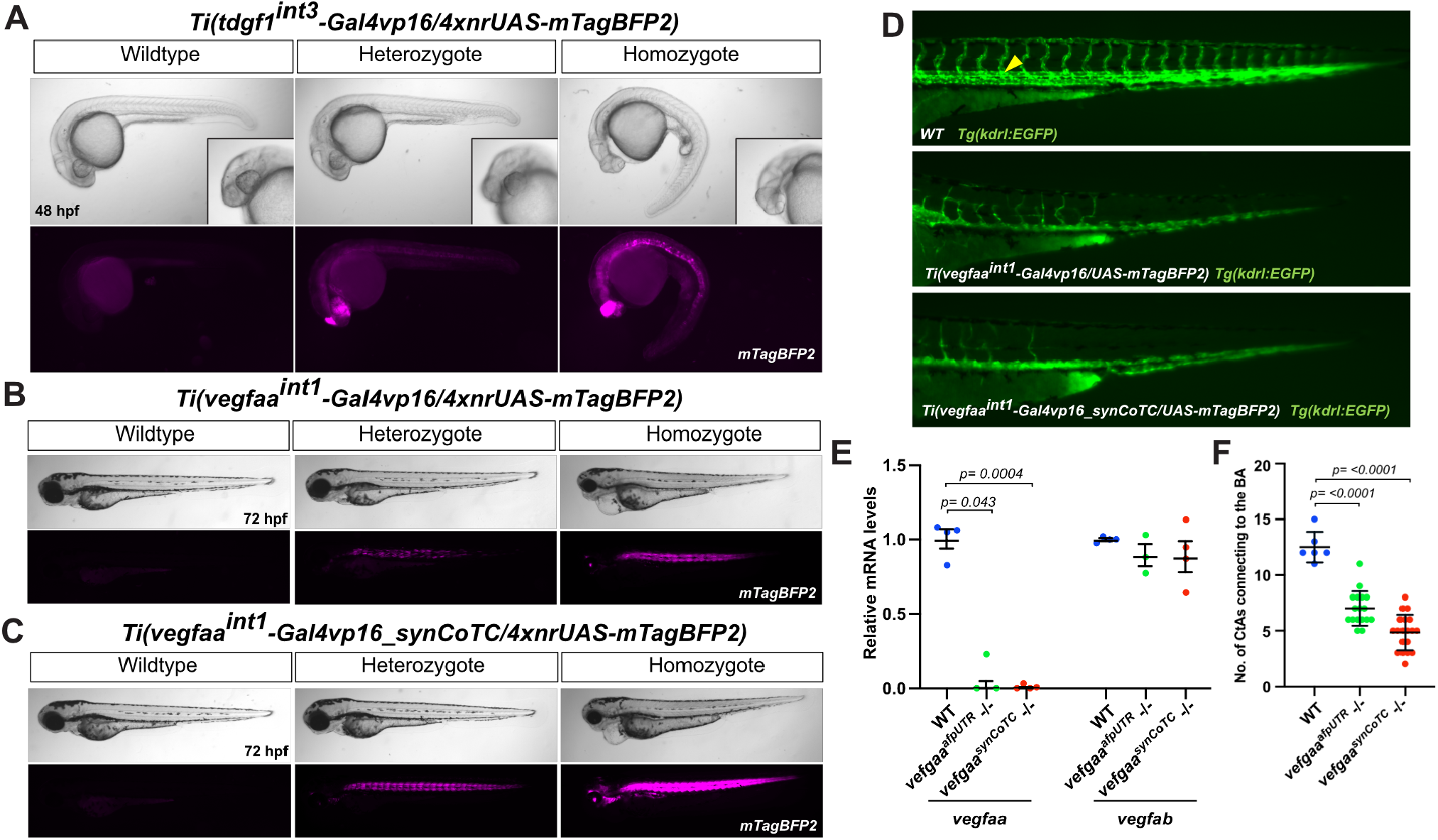
Site-specfic intergration of CRIMPkit vectors into *tdgf1* and *vegfaa*. (**A**) *Ti(tdgf1*^*int3*^*-Gal4vp16/4xnrUAS-mTagBFP2)* targeted insertion line demonstrating loss of *tdgf1* phenotypes in homozygous mutants at 48 hpf. *mTagBFP2* fluorescence is false coloured magenta. (**B**) *Ti(vegfaa*^*int1*^*-Gal4vp16/4xnrUAS-mTagBFP2)* transgenic insertion line (*vegfaa*^*afpUTR*^). (**C**) *Ti(vegfaa*^*int1*^*-Gal4vp16_synCoTC/4xnrUAS-mTagBFP2)* transgenic insertion line (*vegfaa*^*synCoTC*^). *mTagBFP2* fluorescence (magenta) in B and C is captured with the same imaging settings, demonstrating more robust expression in the synCoTC containing line. (**D**) Vascular labelling by the *Tg(kdrl:EGFP)* reporter demonstrates an absence of the dorsal aorta (yellow arrow in wildtype) in mutants from both CRIMP lines, as well as branching defects of the trunk intersegmental vessels at 72 hpf. (**E**) Both *vegfaa*^*afpUTR*^ and *vegfaa*^*synCoTC*^ CRIMP *vegfaa* mutants do not undergo genetic compensation. mRNA expression analysis of *vegfaa* and *vegfab* in *vegfaa* CRIMP mutants demonstrates a complete loss of native *vegfaa* transcript, without a change in *vegfab* mRNA levels, demonstrating genetic compensation has not been induced. Error bars represent SEM for at least three independent biological replicates each consisting of a pooled sample of 17–18 embryos. *rpl13* was used as the reference gene. (**F**) *vegfaa*^*afpUTR*^ and *vegfaa*^*synCoTC*^ CRIMP *vegfaa* mutants have a reduction in the number of branching intracerebral central arteries (CtAs) similar to the reported the non-compensating *vegfaa* promoterless mutant. *vegfaa*^*afpUTR-/-*^ n = 17, and *vegfaa*^*synCoTC-/-*^ n = 19, wildtype n = 6. Error bars represent SEM.

As a second example, we targeted *vegfaa*, a gene involved in vasculature development, as this gene has a well-defined mutant phenotype and has been shown to undergo genetic compensation (Rossi et al., 2015; El-Brolosy et al., 2019). We inserted the *pSA0-T2A-Gal4vp16/4xnrUAS-mTagBFP2* vector into intron-1 of *vegfaa*, to generate the *Ti(vegfaa*^*int1*^*-Gal4vp16/4xnrUAS-mTagBFP2)* targeted insertion line (Figure 1B). Similar to *tdgf1*, 96% of injected embryos had detectable mosaic *mTagBFP2* expression, including one embryo displaying the correct expression pattern in the ventral half of the body plan (0.6% of injected embryos). This embryo was raised and successfully passed on the targeted transgenic insertion (Supplementary Figure S5). Heterozygous *Ti(vegfaa*^*int1*^*-Gal4vp16/4xnrUAS-mTagBFP2)* embryos were phenotypically wildtype and homozygous embryos displayed blood pooling, pericardial edema, and no circulation (Figure 1B), phenocopying the published *vegfaa* mutants (El-Brolosy et al., 2019). We noticed that *mTagBFP2* expression was often variable in heterozygous carriers (Supplementary Fig S6). We reasoned that reporter variation might be due to incomplete transcriptional termination of the *Gal4vp16* interfering with the downstream UAS expression cassette, as poly-A signals are known to be poor transcriptional terminators allowing read-through of the RNA polymerase II (Nojima et al., 2013). Therefore, we incorporated a modified Co-Transcriptional Cleavage (CoTC) type terminator element (Nojima et al., 2013) termed synCoTC, after the afp-UTR poly-A signal of *Gal4vp16*. We generated a second *vegfaa* insertion line with the *pSA0-T2A-Gal4vp16_synCoTC/4xnrUAS-mTagBFP2* vector (Figure 1C, hereafter referred to as *vegfaa*^*synCoTC*^). 95% of injected embryos had detectable mosaic *mTagBFP2* expression, and six embryos displaying the highest level of mosaicism were raised to adulthood and screened, three of which passed on the insertion to progeny (50%)(Supplementary Figure S5). There was no difference in the homozygous mutant phenotype of the *vegfaa*^*synCoTC*^ line compared to the no-terminator line (*vegfaa*^*afpUTR*^); however, the *mTagBFP2* expression was much brighter and more robust, without variation in the *mTagBFP2* levels (Figure 1C), demonstrating that inclusion of the synCoTC terminator prevents variable expression and produces insertional lines with higher levels of reporter fluorescence. Therefore, we incorporated the synCoTC into all vectors in the provided toolkit. *vegfaa* mutants are reported to have vascular defects; therefore, we investigated the vascular system in both our *vegfaa* mutants using the *Tg(kdrl:EGFP)* reporter line. Both CRIMP mutant lines lacked a dorsal aorta and displayed a disruption of intersegmental vessels branching in the tail (Figure 1D), consistent with published *vegfaa* mutants (Rossi et al., 2016), further demonstrating the CRIMPkit vectors are capable of generating null mutant alleles.

Premature stop mutations prior to the last exon in *vegfaa* have been shown to induce genetic compensation, resulting in an upregulation of the orthologous gene *vegfab* (Rossi et al., 2015; El-Brolosy et al., 2019). The non-compensating promoter deletion mutant of vegfaa displays a stronger intracerebral central arteries (CtAs) branching defect than *vegfaa* compensating mutants (El-Brolosy et al., 2019). qRT-PCR analysis in our CRIMP mutants identified a loss of *vegfaa* mRNA without upregulation of *vegfab* (Figure 1E). Quantification of CtA branching in our CRIMP mutants identified a reduction in the branching of the CtAs (Figure 1F) at a similar level to that of the non-compensating promoter-less *vegfaa* mutants (El-Brolosy et al., 2019). Together these results demonstrate that CRIMP insertional mutants do not undergo genetic compensation.

We are interested in high-throughput drug screening and the ability to visually select homozygous mutants to increase throughput and reduce costs. To enable this we incorporated the splitGFP (Kamiyama et al., 2016) system into CRIMPkit vectors. We inserted the *pSA0-mTagBFP2-T2A-splitGFP1-10* vector into intron-2 of *actc1b*, to generate the *Ti(actc1b*^*int2*^*-mTagBFP2-T2A-splitGFP1-10)* transgenic insertion line (Figure 2A, S7). A second allele was generated by inserting the *pSA0-mTagBFP2-T2A-splitGFP11×7* vector into intron-4 of *actc1b*, to generate the *Ti(actc1b*^*int4*^*-mTagBFP2-T2A-splitGFP11×7)* transgenic insertion line (Figure 2B, S7). Crossing these two lines together results in heterozygous embryos that are *mTagBFP2* positive but negative for *GFP* fluorescence. Homozygous embryos (or compound heterozygotes) can be identified by positive *GFP* fluorescence (Figure 2C).

**Figure 2.**
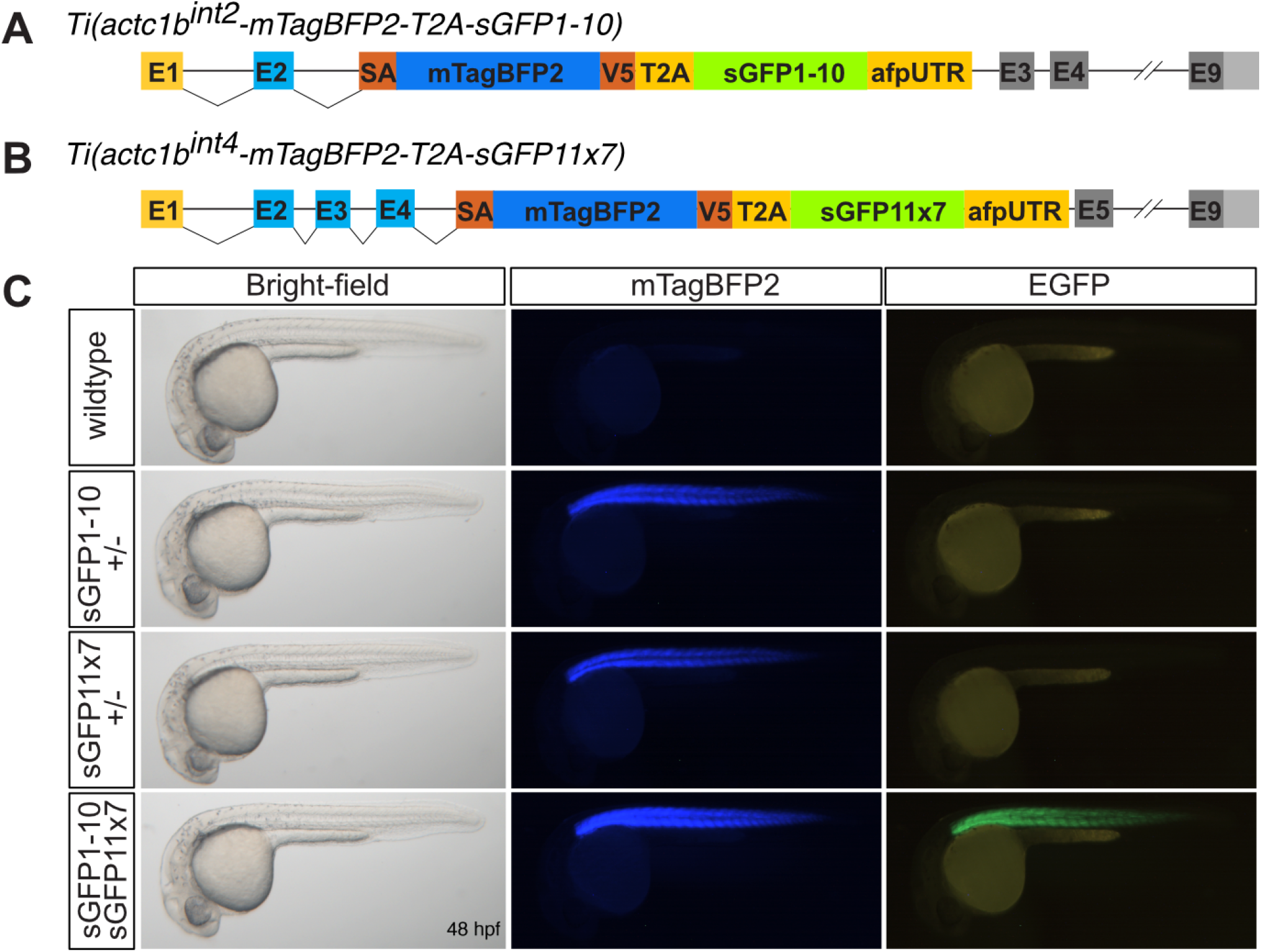
SplitGFP alleles enable visual selection of homozygous mutants. (**A**) Targeted insertion of *pSA_0_mTagBFP2-T2A-sGFP1-10* into *actc1b* Intron-2 to generate the *Ti(actc1b*^*int2*^*-mTagBFP2-T2A-sGFP1-10)* line. (**B**) Targeted insertion of *pSA_0_mTagBFP2-T2A-sGFP11×7* into *actc1b* Intron-4 to generate the *Ti(actc1b*^*int4*^*-mTagBFP2-T2A-sGFP11×7)* line. (**C**) Embryos were generated by crossing the two sGFP lines together. Heterozygous carriers display only *mTagBFP2* expression, whereas compound heterozygous mutants express both *mTagBFP2* and *EGFP*, allowing visual determination of genotype.

The CRIMP system provides a significant advantage over current strategies for multiple reasons. Firstly, the CRIMPkit vectors are universal as they are available in all three reading frames after the splice acceptor, enabling this toolkit to target any intron of any target gene. Secondly, CRIMPkit vectors are ready-to-use without modification of the targeting vectors prior to use, with options for both high and low expressed genes (with and without *Gal4/UAS* fluorophore amplification), and multiple fluorophore reporters (*mTagBFP2, mKate2*, and *splitGFP*). Thirdly, the high efficiency of this system and early timing of integration events can enable efficient pre-selection of likely founders in F_0_ embryos after injection, reducing the time and effort required for screening. Finally, CRIMP alleles can produce a complete loss of function, terminating transcription at the insert site, without the induction of genetic compensation. Whilst the provided toolkit is optimised for use in zebrafish, due to codon optimisation of the plasmids, the approach and toolkit could be used in any species.

## Methods

### CRIMPkit vector availability

CRIMPkit vectors are available from Addgene as a plasmid collection: https://www.addgene.org/kits/bryson-richardson-crimpkit/.

### CRIMPkit vector design

The vector design contains a highly active guideRNA binding site (Hbait; that is not present in the zebrafish genome) for CRISPR/Cas9 mediated linearisation. The Hbait site is directly upstream of the splicing cassette which consists of the 3’ region of *β-actin* intron-2, including the splice acceptor for exon-3. One or two bases have been added following the splice acceptor in pSA_1 and pSA_2 vectors respectively, to ensure the downstream reading frame is preserved, a flexible protein linker (3xGGGGS), a fluorophore (*mTagBFP2* or *mKate2*) or *Gal4vp16* and a UAS fluorophore cassette, a 3’UTR and poly-A signal derived from ocean pout antifreeze protein 3’UTR (afpUTR) (Horstick et al., 2015; Gibbs and Schmale, 2000), followed by a Co-transcriptional Cleavage (CoTC)-type terminator element (to prevent read-through of the RNA polymerase II) termed synCoTC. The synCoTC consists of the afpUTR including the poly-A signal, followed by the human CCNB1 CoTC (Nojima et al., 2013).

The transcriptional activator Gal4vp16 was selected to maximise fluorophore expression levels in targeted genes with low expression levels, where using a direct fluorophore expression may not produce sufficient levels for visual identification. The 4xnrUAS element used in the Gal4/UAS vectors is methylation resistant to prevent silencing in subsequent generations (Akitake et al., 2011). The 4xnrUAS cassettes include the modified UBC-intron before the *mTagBFP2* or *mKate2* coding sequences to increase expression levels (Horstick et al., 2015).

For the fluorophores, *splitGFP* template sequences were from taken Kamiyama et al. (2016). *mTagBFP2* (Subach et al., 2011) and *mKate2* (Shcherbo et al., 2009)) under control of the 4xnrUAS has an additional valine at the second position for increased mRNA stability and expression levels (Balleza et al., 2018). This enhancement is present in some of the non-Gal4 vectors as detailed in (refer to Table S1 for plasmid details). In addition to the splice incorporating vectors, the CRIMPkit includes Kozak vectors, which lack a splicing cassette, for targeting 5’UTR regions.

The CRIMPkit vectors have been codon optimised for zebrafish using the CodonZ software to enhance expression levels (Horstick et al., 2015).

In the CRIMPkit vectors the CRISPR/Cas9 Hbait guide site is flanked by 48bp FRT3 and FRT sites, which allow for recombination-mediated cassette exchange (RMCE) that can be induced using the *FLP* protein (Turan et al., 2011). This is untested, but is expected to allow for exchange of the CRIMPkit insertion with alternative sequences if desired.

A number of vectors include the V5 peptide tag (GKPIPNPLLGLDST) (Hanke et al., 1992). The Gal4/UAS vectors have a V5 peptide tag included directly after the splice acceptor, before the T2A-Gal4 element, which enables the truncated protein, if produced, to be detected via immunohistochemistry or western blot using a antibody against V5. The *pSA_X_mTagBFP2_synCoTC* vectors also have a flexible linker (GGGGS)-V5 tag at the C-terminal of *mTagBFP2*. All CRIMPkit vectors that contain *mTagBFP2-T2A-splitGFP* have the GGGGS linker-V5 tag on the C-terminal of *mTagBFP2* before the T2A-splitGFP.

### CRIMP Protocol

#### G2 cell cycle phase and Cas9 protein

In medaka it has been reported that targeted-integration only occurs during the G2 phase of the cell cycle (Gutierrez-Triana et al., 2018), and we suspected this would be similar in zebrafish. Zebrafish have a single G2-phase during the first cell division, after which G2 does not occur again until around the midblastula-transition (Bouldin and Kimelman, 2014). We reasoned that the lag in Cas9 production when injecting Cas9 mRNA would miss this early G2 phase and result in lower integration efficiency. In order to facilitate early integration events during this first cell division we used pre-complexed guideRNA (Alt-R™), Cas9 protein (Cas9 HiFi v3 from Integrated DNA Technologies), and targeting plasmid in our injection mix.

#### guideRNA concentration and type

guideRNA molecules are well tolerated even when injected at high concentrations, however, the contaminates resulting from the purification of in-house synthesised guideRNA’s, often cause toxicity when injected at high concentrations (Shah et al., 2015). We suggest using the two-part Alt-R™ crRNA and tracrRNA system (Integrated DNA Technologies) for guideRNA’s as these have high efficiency cutting, are resistant to RNase degradation and, in our experience, do not cause toxicity when injected at high concentrations (we regularly inject them at a final concentration of >300 ng/μl without toxicity).

#### guideRNA design and location

Intronic guideRNA sites are identified using the IDT™ online tool (https://sg.idtdna.com/site/order/designtool/index/CRISPR_CUSTOM), by manually entering intronic sequences and selecting *Danio rerio* as the species. Target sites with the highest on-target and off-target scores are selected, generally with a preference to those located in the 3’ half of the intron (although this is not mandatory). guideRNA target sites used in this study are listed in Table S2.

#### Ribonucleoprotein complex solubility

KCL salt was included in the injection mix to increase the solubility of the Cas9 ribonucleoprotein complex and increase the likelihood of early targeting, as per Burger et al. (2016).

### Fish Maintenance

Fish were maintained in the Monash University Fish Core facility under breeding colony licenses MARP/2015/004/BC and ERM22161. The creation of transgenic lines was approved by the School of Monash University Animal Ethics Committees (ERM18912, 21168). All experiments were carried out on embryos of TU/TL background. All animal work was approved by the Monash Animal Ethics Committee in accordance with the care and use of animals for scientific purposes. Fish were anaesthetised for live imaging using Tricaine methanesulfonate (3-amino benzoic acidethylester; Sigma Aldrich, E10521) at a final concentration of 0.016% in E3 embryo medium (5 mM NaCl, 0.17 mM KCl, 0.33 mM CaCl2, 0.33 mM MgSO4, 0.00004% [v/v] methylene blue in water, pH 7.2).

### cDNA synthesis and qPCR

Total RNA was extracted using TRIzol^™^ reagent (Thermo Fisher Scientific). cDNA was synthesized from 1μg of each RNA sample in a 20 μl reaction using ProtoScript^®^ II First Strand cDNA synthesis kit (New England Biosciences) and oligo(dT)20 and random hexamer primers following the supplier’s instructions. Quantitative PCR (qPCR) was performed on a LightCycler^®^ 480 instrument using LightCycler^®^ 480 SYBR^®^ Green I Master mix (Roche), as per manufacturers protocol. *vegfaa* samples were normalized against *rpl13* (El-Brolosy et al., 2019) as a reference gene. Primers for quantitative PCR are listed in Table S2.

### Imaging

Live Imaging was performed with an Olympus SZX16 stereomicroscope. *mTagBFP2* and *mKate2* fluorescence was visualised using the Chroma™ 49021 and 49008 filter sets respectively. In order to examine intracerebral central arteries in the brain at 60 hpf, embryos were fixed in 4% paraformaldehyde overnight at 4°C, washed in PBST (1× PBS, 0.1% Tween 20 (Sigma)), and set in 1% low melting agarose in 0.8 mm fluorinated ethylene propylene (FEP) tubing (Bola, Grünsfeld, Germany). Genotypes were randomised, and the investigator blinded to genotype. Images were taken using a Thorlabs confocal microscope (Newton, NJ, USA), with an Olympus 20x water dipping 1.0 NA objective (Tokyo, Japan), pinhole 25 μm, 2.005 μm/pixel, step size 1 μm, averaging 16 frames. Images were analysed using Fiji (ImageJ) software (Schindelin et al., 2012).

### Microinjection

Freshly fertilised embryos were injected using a Femtojet microinjector (Eppendorf). Approximately ∼1 nl of the mix was injected into the cell portion of the embryo at the one-cell stage, usually within the first 10 minutes of being laid.

## Data availability

Raw data can be obtained from the Bridges repository: https://bridges.monash.edu/projects/CRIMP_A_CRISPR_-Cas9_Insertional_Mutagenesis_Protocol_and_the_CRIMP_Toolkit/171171

