## Supplementary Material for "CRIMP: A CRISPR/Cas9 Insertional Mutagenesis Protocol and the CRIMP Toolkit"

### Figure S1: Schematic of CRIMP vector design

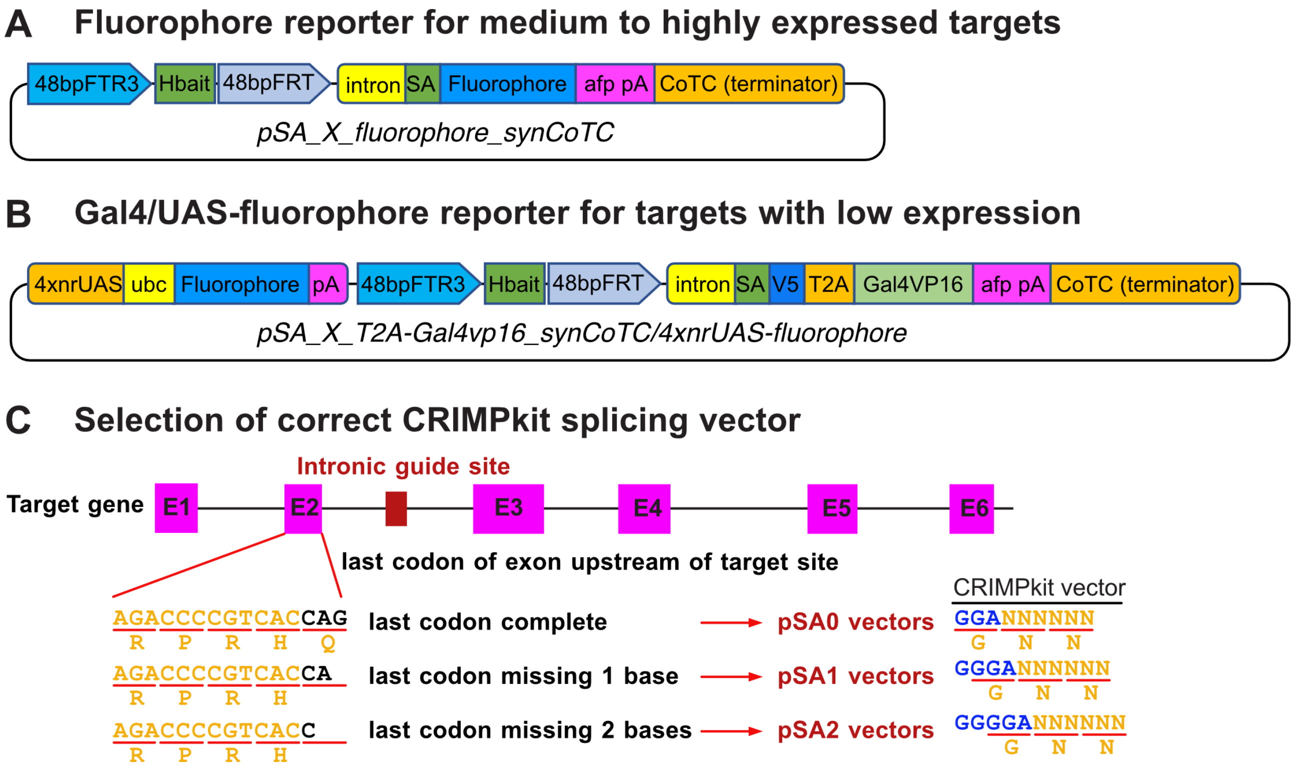


**Figure S1. Schematic of CRIMP vector design**. CRIMP vectors contain a guideRNA target site (Hbait) for CRISPR/Cas9 mediated linearisation. The Hbait site is directly upstream of the splicing cassette which consists of the 3' region of $\beta$-*actin* intron-2, including the splice acceptor for exon-3. One or two bases have been added following the splice acceptor in pSA_1 and pSA_2 vectors respectively, to ensure the downstream reading frame is preserved. CRIMPkit vectors with a fluorophore after the splice acceptor (**A**) are designed for targeting genes with high expression levels. Vectors with Gal4vp16 after the splice acceptor and downstream UAS:fluorophore cassette (**B**) are for targeting genes with low expression levels. SA = Splice Acceptor, afp pA = Ocean pout anti-freeze protein 3'UTR until the poly-A sequence. ubc = 5'UTR intron of ubiquitin C, CoTC = Co-Transcriptional Cleavage. (**C**) Selection of correct the CRIMPkit vector. The reading frame of the last codon in the exon upstream of the intronic target site will determine the correct vector to use in order to preserve the reading frame of the downstream coding sequence in a successfully targeted intron. A complete final codon will require a pSA0- vector, a codon missing one base requires a pSA1- vector, and a codon missing two bases requires a pSA2- vector. Black bases indicate the final codon of an exon upstream of an intronic target. Blue bases indicate the sequence incorporated in CRIMPkit vectors to maintain reading frame after splicing.

#### Table S1. List of toolkit vectors

| CRIMP plasmid name | Addgene number |
|---|---|
| pSA_0_mKate2_synCoTC | 199469 |
| pSA_1_mKate2_synCoTC | 199470 |
| pSA_2_mKate2_synCoTC | 199471 |
| pSA_0_mTagBFP2_synCoTC | 199472 |
| pSA_1_mTagBFP2_synCoTC | 199473 |
| pSA_2_mTagBFP2_synCoTC | 199474 |
| pSA_0_mTagBFP2-T2A-sGFP1-10_synCoTC | 199475 |
| pSA_1_mTagBFP2-T2A-sGFP1-10_synCoTC | 199476 |
| pSA_2_mTagBFP2-T2A-sGFP1-10_synCoTC | 199477 |
| pSA_0_mTagBFP2-T2A-sGFP11x7_synCoTC | 199478 |
| pSA_1_mTagBFP2-T2A-sGFP11x7_synCoTC | 199479 |
| pSA_2_mTagBFP2-T2A-sGFP11x7_synCoTC | 199480 |
| pSA_0_T2A-Gal4vp16_synCoTC/4xnrUAS-mKate2 | 199481 |
| pSA_1_T2A-Gal4vp16_synCoTC/4xnrUAS-mKate2 | 199482 |
| pSA_2_T2A-Gal4vp16_synCoTC/4xnrUAS-mKate2 | 199483 |
| pSA_0_T2A-Gal4vp16_synCoTC/4xnrUAS-mTagBFP2 | 199484 |
| pSA_1_T2A-Gal4vp16_synCoTC/4xnrUAS-mTagBFP2 | 199485 |
| pSA_2_T2A-Gal4vp16_synCoTC/4xnrUAS-mTagBFP2 | 199486 |
| pSA_0_T2A-Gal4vp16_synCoTC/4xnrUAS-mTagBFP2-T2A-sGFP1-10 | 199487 |
| pSA_1_T2A-Gal4vp16_synCoTC/4xnrUAS-mTagBFP2-T2A-sGFP1-10 | 199488 |
| pSA_2_T2A-Gal4vp16_synCoTC/4xnrUAS-mTagBFP2-T2A-sGFP1-10 | 199489 |
| pSA_0_T2A-Gal4vp16_synCoTC/4xnrUAS-mTagBFP2-T2A-sGFP11x7 | 199490 |
| pSA_1_T2A-Gal4vp16_synCoTC/4xnrUAS-mTagBFP2-T2A-sGFP11x7 | 199491 |
| pSA_2_T2A-Gal4vp16_synCoTC/4xnrUAS-mTagBFP2-T2A-sGFP11x7 | 199492 |
| pKozak-mTagBFP2_synCoTC | 199493 |
| pKozak-mTagBFP2-T2A-sGFP1-10_synCoTC | 199494 |
| pKozak-mTagBFP2-T2A-sGFP11x7_synCoTC | 199495 |
| pKozak-Gal4vp16_synCoTC/4xnrUAS-mTagBFP2 | 199496 |
| pKozak-Gal4vp16_synCoTC/4xnrUAS-mTagBFP2-T2A-sGFP1-10 | 199497 |
| pKozak-Gal4vp16_synCoTC/4xnrUAS-mTagBFP2-T2A-sGFP11x7 | 199498 |

### Figure S2: CRIMP Injection protocol

#### 1 Resuspend Alt-R crRNA and tracrRNA in Nuclease-Free IDTE Buffer to final stock concentrations of 100 µM each

- 2 nM = 20 µl
- 5 nM = 50 µl

This is stored in the -20°C freezer once resuspended

---

#### 2 Make guideRNA complex

Make fresh gRNA complexes for each injection

**Note: you will need 1 µl genomic gudieRNA and 0.5 µl Hbait (plasmid) guideRNA per injection**

2.1 Mix the following components to create a 30 µM gRNA solution: - only make enough for your injection.

| Component | Pick the most suitable volumes for your requirements | |
|---|---|---|
| 100 µM Alt-R™ CRISPR-Cas9 crRNA | 0.25 µl | 0.4 µl |
| 100 µM Alt-R™ CRISPR-Cas9 tracrRNA | 0.25 µl | 0.4 µl |
| Nuclease-Free Duplex Buffer (IDT) | 0.33 µl | 0.53 µl |
| Final volume | 0.83 µl | 1.33 µl |

2.2 Heat at 95°C for 5 min (in PCR machine).

2.3 Remove from heat, and allow to cool to room temperature (15–25°C) on your bench top.

Note: the final concentration for the crRNA is 360 ng/µl and for the tracrRNA is 670 ng/µl.

Therefore total guideRNA complex concentration is 1030 ng/µl (30 µM = 1030 ng/µl).

**Note2: this complex can be stored at -20°C for up to three months with no loss of activity (when at least 30 µM), but is best to make fresh.**

#### 3 Assemble the mix (CRISPR/Cas9 complexes) for injection:

| Component | Amount | Final concentration |
|---|---|---|
| 1M KCL (0.2 µm filter sterilised) [1] | 1.5 µl | 300 mM |
| genomic guideRNA complex (from step 2.3) @1030 ng/µl | 1µl | 206 ng/µl (pg/µl)[2] |
| Hbait plasmid guideRNA complex (from step 2.3) @1030 ng/µl | 0.5 µl | 103 ng/µl (pg/nl)[2] |
| Targetting plasmid -> 36 fmol total plasmid [3] (dilute so 0.5 µl is sufficient) | 0.5 µl | 7.2 fmol/µl |
| phenol red (0.05%) (0.2 µm filter sterilised) | 0.5 µl | 0.005% |
| H20 (sterile MilliQ) | to final volume 5µl | |
| Cas9 HiFi V3 protein (IDT™) (62 µM = 10,000 ng/µl) | 0.3 µl | 700 ng/µl (pg/nl) |
| Final volume | 5µl | |

[1] Required for solubility of complex - see Burger et al. (2016)
[2] See Shah et al. (2015) describing non-lethal injection of high amounts of guideRNA
[3] Calculate plasmid fmoles at NebBioCalculator https://nebiocalculator.neb.com/#!/dsdnaamt

---

#### 4 Incubate at 37°C for 10 min - 1 hour before injecting into one-cell stage embryos

---

#### 5 Inject embryos with CRISRP/Cas9 Ribonuclear (RNP) complex)

5.1 Collect embryos within 5 minutes of being laid and inject 2 nL of CRISRP/Cas9 Ribonuclear (RNP) complex directly into the cell of the embryo.

Note: In our experience injecting into the embryos withing the first 15 minutes increases the likelihood of integration before (or during) the first cell division, and maximised the change of obtaining embryos with reporter expression in half of the embryo body plan.

5.2 Incubate injected embryos at 28 °C and remove any unfertilised or dead embryos before leaving for the day.

5.3 Screen injected embryos for positive reporter expression at 24 hpf (or a time point after target gene is expressed).

---

#### Figure S3. Example of successful targeted integration during the first cell division

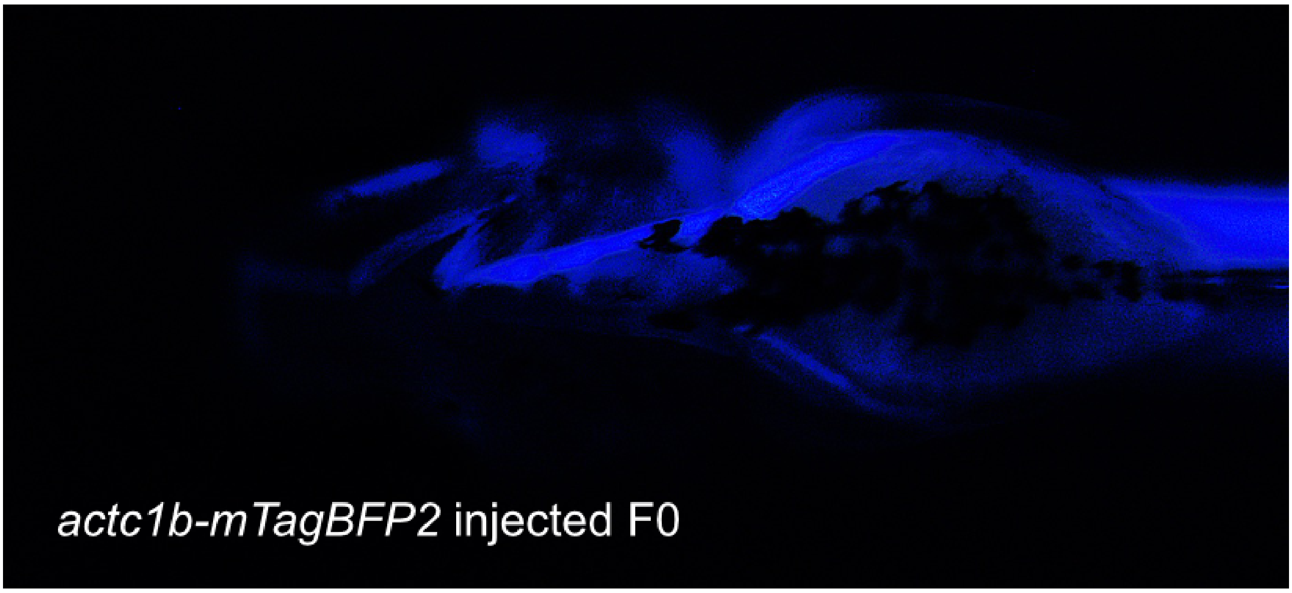


**Figure S3. Example of successful targeted integration during the first cell division.** Integration of the targeting vector into intron-2 of *actc1b* during the first cell division results in embryos with *mTagBFP2* expression in one half of the ventral body plan. In our experience, such embryos, when raised to adulthood and crossed, have always transmitted the successful integration event to their progeny. Ventral view of a 4 dpf embryo.

#### Figure S4. location and integration site for *Ti(tdgf1$^{int3}$-Gal4vp16/4xnrUAS-mTagBFP2)* transgenic line

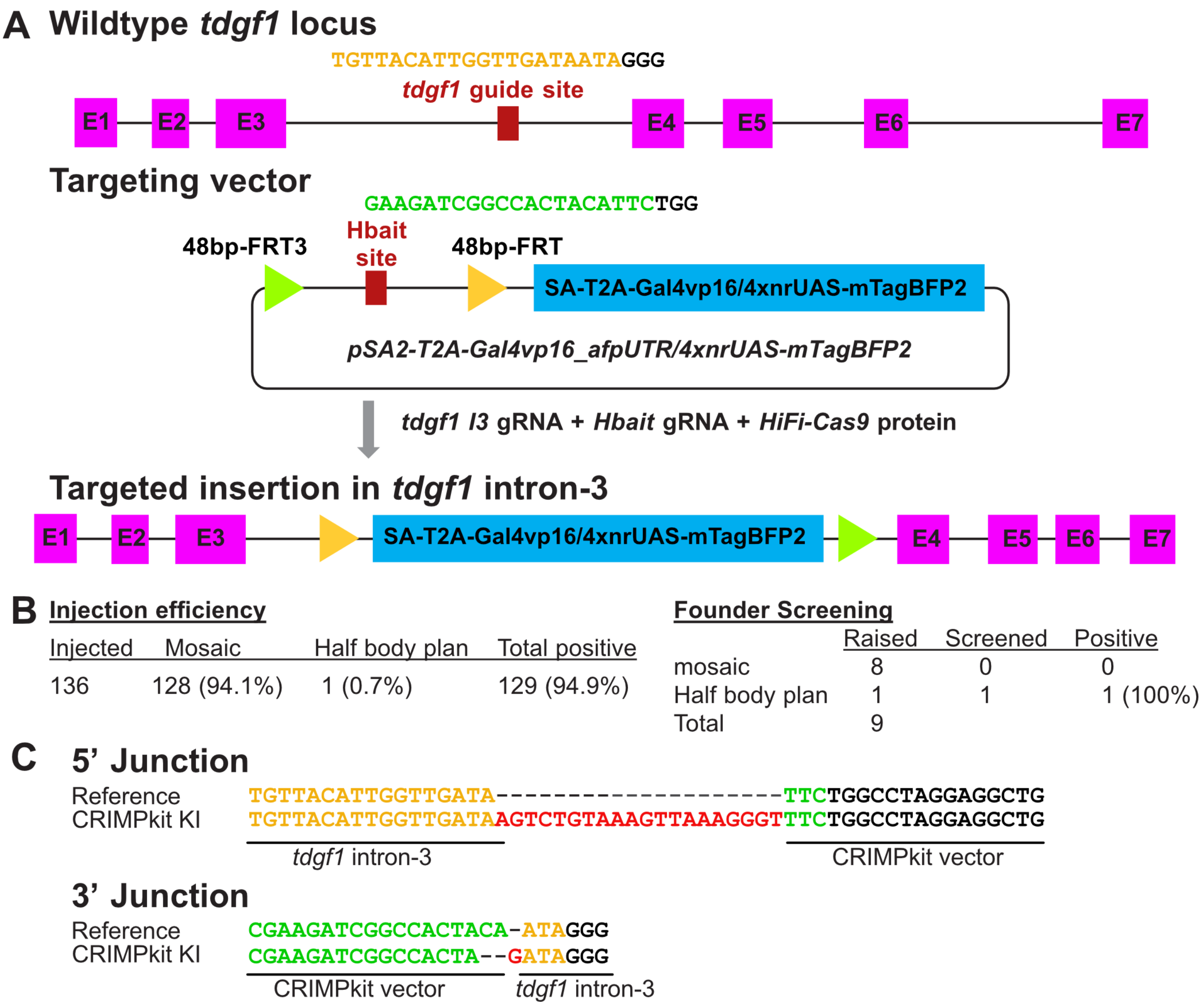


**Figure S4. location and integration site for *Ti(tdgf1$^{int3}$-Gal4vp16/4xnrUAS-mTagBFP2)* transgenic line.** (**A**) Targeting strategy for *tdgf1* insertional mutant. The *tdgf1* guideRNA site is located in intron-3. The *pSA2-T2A-Gal4vp16_-afpUTR/4xnrUAS-mTagBFP2* targeting vector was inserted into the intron-3 target site. (**B**) Injection and founder screening efficiencies. (**C**) Sequences of the 5' and 3' insertion sites. *tdgf1* guideRNA sequence is coloured yellow, Hbait guideRNA sequence coloured green. Red indicates extra bases added during insertion.

### Figure S5. location and integration sites for *vegfaa$^{afpUTR}$* and *vegfaa$^{synCoTC}$* transgenic lines

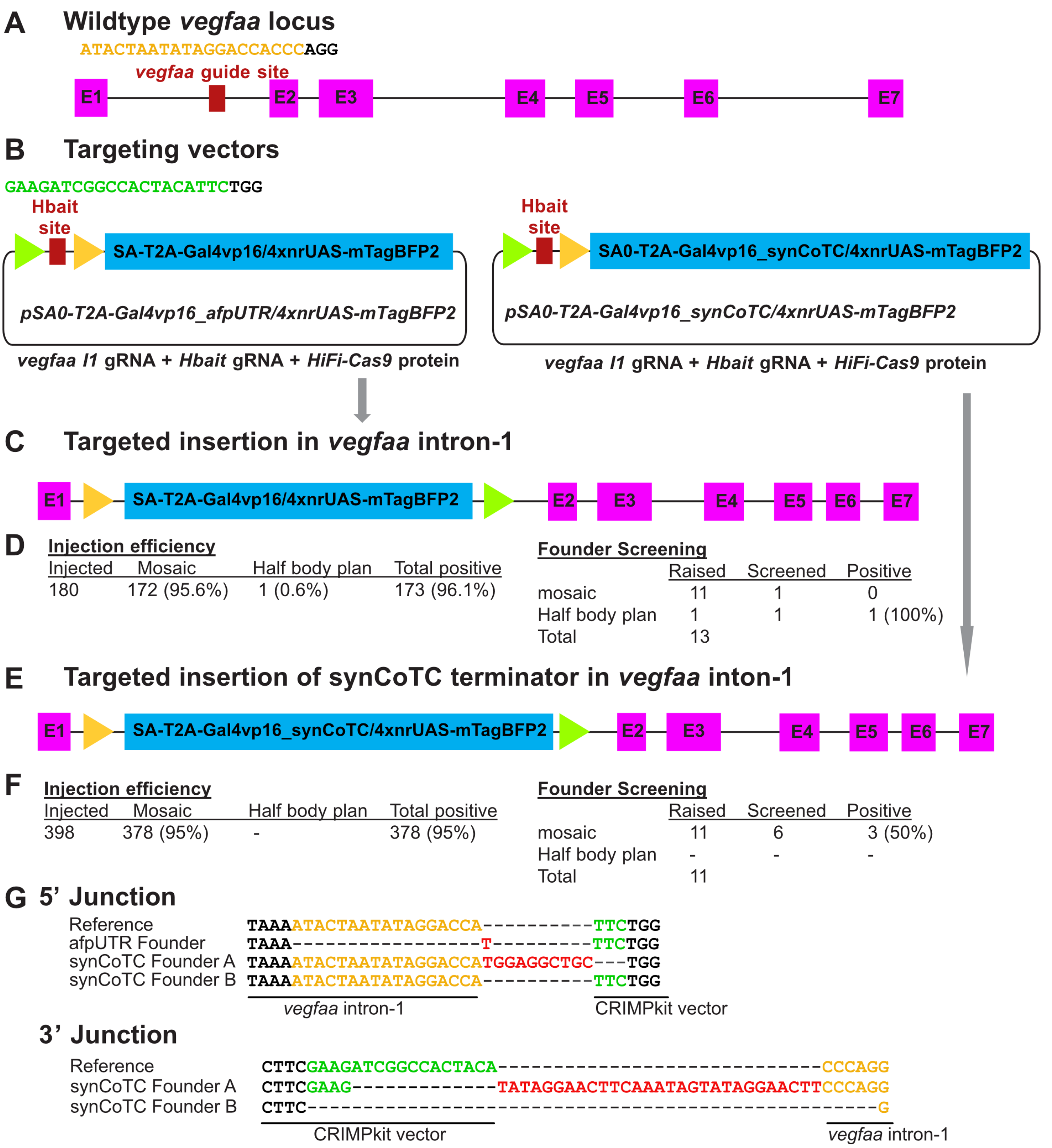


**Figure S5. location and integration site for *Ti(vegfaa$^{int1}$-Gal4vp16/4xnrUAS-mTagBFP2)* and *Ti(vegfaa$^{int1}$-Gal4vp16_synCoTC/4xnrUAS-mTagBFP2)* transgenic lines**. (**A**) Targeting strategy for *vegfaa* insertional mutants. The *vegfaa* guideRNA site is located in intron-1. (**B**) The afpUTR only targeting vector *pSA0-T2A-Gal4vp16_afpUTR/4xnrUAS-mTagBFP2* and the synCoTC terminator containing vector *pSA0-T2A-Gal4vp16_synCOTC/4xnrUAS-mTagBFP2* were targeted to *vegfaa* intron-1 site to generate the (**C**) *Ti(vegfaa$^{int1}$-Gal4vp16/4xnrUAS-mTagBFP2)* line and (**E**) *Ti(vegfaa$^{int1}$-Gal4vp16_synCoTC/4xnrUAS-mTagBFP2)* lines respectively. Injection and founder screening efficiencies of both lines are shown in (**D**) and (**F**). (**G**) Sequences of the 5' and 3' insertion sites. *vegfaa* guideRNA sequence is coloured yellow, Hbait guideRNA sequence coloured green. Red indicates extra bases added during insertion.

#### Table S2. Primer and guideRNA sequences

| Genotyping primers | Primer sequence |
|---|---|
| *tdgf1_int3_F* | TCTCTGGGAATGTCATGGCT |
| *tdgf1_int3_R* | AGGTGTCTAGCATTGCGGTA |
| *$\beta$-intron_R* | ACCAGCTCACCGAGAAATGA |
| *mTagBFP_end_F* | AGCTGGGACACAAGCTGAAT |
| *vegfaa_int2_F2* | TCTGTAGGCGGGAAAGAAGA |
| *vegfaa_int2_R4* | AGCCAAATATTGTCCTAACAAAGTG |
| *T2A_gal4_qPCR_R* | GAAGTTTCATTGGGCCAGGG |
| qRT-PCR primers | |
| *vegfaa_qPCR_F* | CCCACGATATCACACTCGGT |
| *vegfaa_qPCR_R* | GGATGTACGTGTGCTCGATCT |
| *vegfab_qPCR_F* | GGTGCTGCAATGATGAAATG |
| *vegfab_qPCR_R* | TGTCACCCTGATGACGAAGA |
| *rpl13_qPCR_F* | TAAGGACGGAGTGAACAACCA |
| *rpl13_qPCR_R* | CTTACGTCTGCGGATCTTTCTG |
| crRNA guide sequence(PAM) | |
| *tdgf1_int3* | TGTTACATTGGTTGATAATA(GGG) |
| *vegfaa_int1* | ATACTAATATAGGACCACCC(AGG) |
| *actc1b_int2* | TAGATTTAGGACAAGTCTGC(TGG) |
| *actc1b_int4* | ATCTACTTAGACCTCACATG(TGG) |
| Hbait | GAAGATCGGCCACTACATTC(TGG) |

#### Figure S6. Reporter expression but not phenotype is variable in the absence of a transcriptional terminator

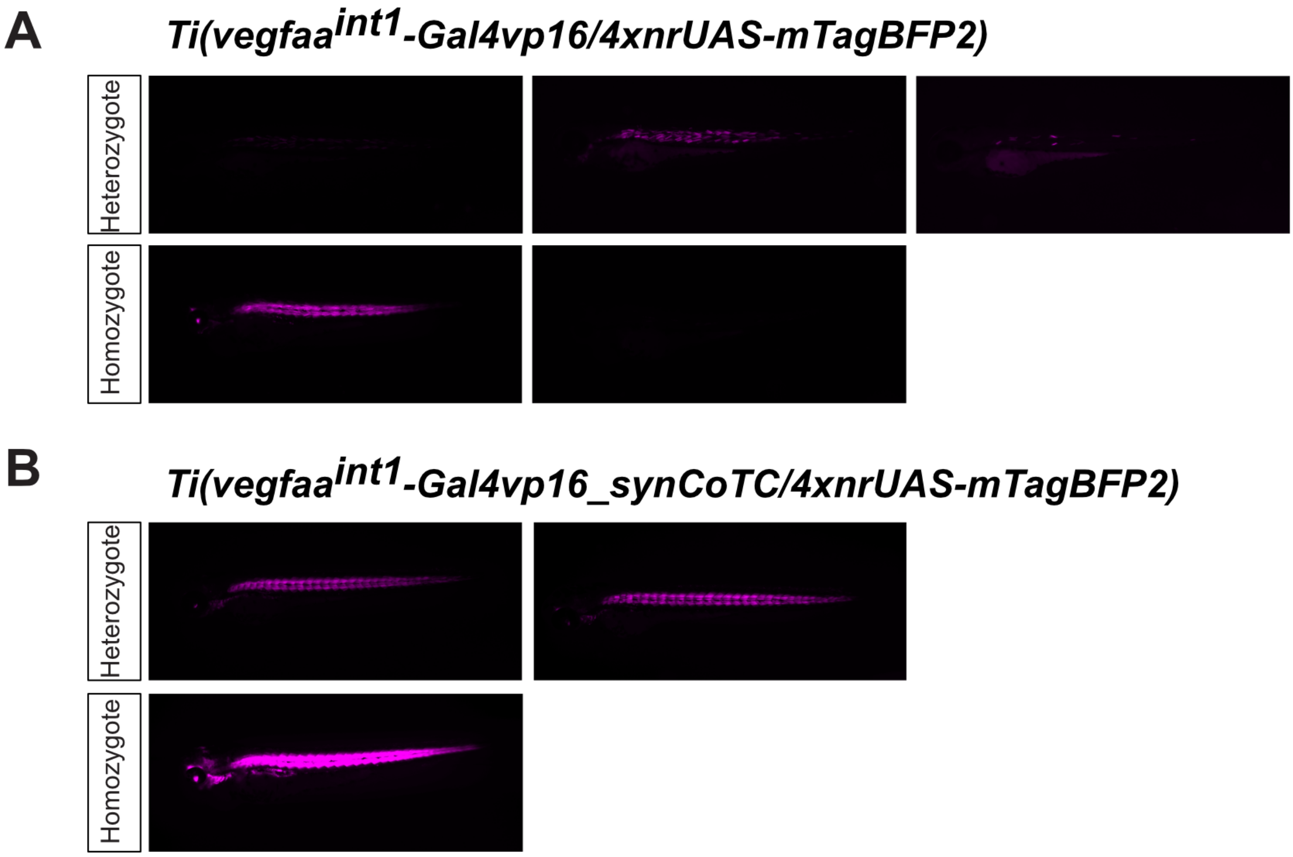


**Figure S6. Reporter expression but not phenotype is variable in the absence of a transcriptional terminator.** (**A**) The *Ti(vegfaa$^{int1}$-Gal4vp16/4xnrUAS-mTagBFP2)* line that lacks a transcriptional terminator displays variation in both the level and pattern of *mTagBFP2* expression (magenta). The mutant phenotype is not affected. (**B**) Inclusion of the synCoTC terminator in the *Tgi(vegfaa$^{int1}$-Gal4vp16_synCoTC/4xnrUAS-mTagBFP2)* line results in robust levels of reporter expression in all embryos without spatial variation. All images are taken at the same exposure.

### Figure S7. Targeting strategy for *Ti(actc1b$^{int2}$-mTagBFP2-T2A-sGFP1-10)* and *Ti(actc1b$^{int4}$-mTagBFP2-T2A-sGFP11x7)* transgenic lines

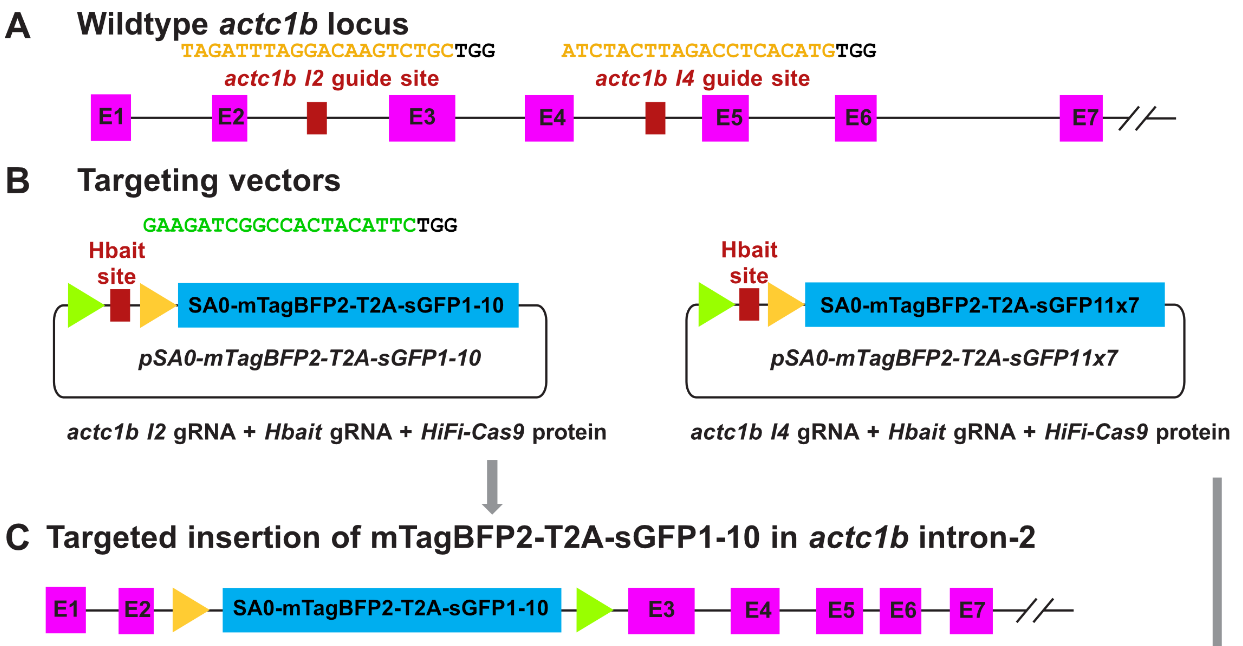


**D**

**Injection efficiency**

| Injected | Mosaic | Half body plan | Total positive |
|---|---|---|---|
| 93 | 12 (12.9%) | 2 (2.2%) | 14 (15.1%) |

**Founder Screening**

| | Raised | Screened | Positive |
|---|---|---|---|
| mosaic | 2 | 2 | 0 |
| Half body plan | 1 | 1 | 1 (100%) |
| Total | 3 | | |

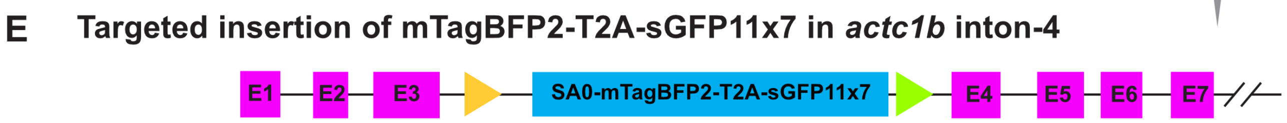


**F**

**Injection efficiency**

| Injected | Mosaic | Half body plan | Total positive |
|---|---|---|---|
| 377 | 42 (11%) | 1 (0.2%) | 43 (11.4%) |

**Founder Screening**

| | Raised | Screened | Positive |
|---|---|---|---|
| mosaic | 5 | 4 | 1 (25%) |
| Half body plan | 1 | 1 | 1 (100%) |
| Total | 6 | | |

**G 5' Junction**

```
Reference (int-4)   ATCTACTTAGACCTCAC------------------------TTCTGG
sGFP11x7            ATCTACTTAGACCTCA----+848 bp plasmid seq---TTCTGG
                    actc1b intron-4                           CRIMPkit vector
```

**Figure S7. Targeting strategy for *Ti(actc1b$^{int2}$-mTagBFP2-T2A-sGFP1-10)* and *Ti(actc1b$^{int4}$-mTagBFP2-T2A-sGFP11x7)* transgenic lines.** (**A**) Target sites for *actc1b* insertional mutants. *actc1b* guideRNA sites are located in intron-2 and intron-4. (**B**) The *pSA0-mTagBFP2-T2A-sGFP1-10* targeting vector and *pSA0-mTagBFP2-T2A-sGFP11x7* vectors were targeted to *actc1b* intron-1 and intron-4 respectively, to generate the (**C**) *Ti(actc1b$^{int2}$-mTagBFP2-T2A-sGFP1-10)* line and (**E**) *Ti(actc1b$^{int4}$-mTagBFP2-T2A-sGFP11x7)* lines respectively. Injection and founder screening efficiencies of both lines are shown in (**D**) and (**F**). (**G**) Sequence of the 5' insertion site for the sGFP11x7 line. Multiple copies of the plasmid in the target site prevented insert site sequencing in the sGFP1-10 line. *actc1b* guideRNA sequence is coloured yellow, Hbait guideRNA sequence coloured green. Red indicates extra bases added during insertion.
